# *Paraburkholderia edwinii* protects *Aspergillus* sp. from phenazines by acting as a toxin sponge

**DOI:** 10.1101/2021.03.28.437412

**Authors:** Kurt M. Dahlstrom, Dianne K. Newman

## Abstract

Many environmentally and clinically important fungi are sensitive to toxic, bacterially-produced, redox-active molecules called phenazines. Despite being vulnerable to phenazine-assault, fungi inhabit microbial communities that contain phenazine producers. Because many fungi cannot withstand phenazine challenge, but some bacterial species can, we hypothesized that bacterial partners may protect fungi in phenazine-replete environments. In the first soil sample we collected, we co-isolated several such physically associated pairings. We discovered the novel species *Paraburkholderia edwinii* and demonstrated it can protect a co-isolated *Aspergillus* species from phenazine-1-carboxylic acid (PCA) by sequestering it, acting as a toxin sponge; in turn, it also gains protection. When challenged with PCA, *P. edwinii* changes its morphology, forming aggregates within the growing fungal colony. Further, the fungal partner triggers *P. edwinii* to sequester PCA and maintains conditions that limit PCA toxicity by promoting an anoxic and highly reducing environment. A mutagenic screen revealed this program depends on the stress-inducible transcriptional repressor HrcA. We show that one relevant stressor in response to PCA challenge is fungal acidification and that acid stress causes *P. edwinii* to behave as though the fungus were present. Finally, we reveal this phenomenon as widespread among *Paraburkholderia* with moderate specificity among bacterial and fungal partners, including plant and human pathogens. Our discovery suggests a common mechanism by which fungi can gain access to phenazine-replete environments, and provides a tractable model system for its study. These results have implications for how rhizosphere microbial communities as well as plant and human infection sites are policed for fungal membership.

## Introduction

The presence or absence of particular fungal species in host-associated microbial communities plays a central role in human and plant health, crop yield, and climate change ^1–3^. However, we lack an understanding of how key fungal species are integrated into these communities in the face of rampant chemical warfare. It has long been known that the soil is home to diverse microbes that produce natural products with antibiotic activity. Important amongst these are phenazines, redox active compounds that can restrict fungal growth and have been shown to be responsible for excluding fungi from agriculturally important microbial communities ^4,5^. A recent metagenomic study revealed that phenazine biosynthesis capacity is widespread in agricultural soils and crop microbiomes ^6^. Given that drier soils are also associated with higher rates of phenazine producers colonizing wheat, this suggests that soil fungi may need to contend with higher concentrations of phenazines as the climate shifts ^7,8^. Paradoxically, many fungi that are sensitive to phenazines are routinely found living in close proximity to phenazine-producing bacteria, including pathogenic fungi in the lungs of cystic fibrosis patients, beneficial and phytopathogenic fungi in the rhizosphere, and in oceanic environments including coral ^9–12^. This pattern of co-habitation indicates there may be a general way fungi are screened for membership in microbial communities that produce phenazines that holds broad relevance. We set out to identify such a putative screening mechanism, a necessary step towards the goal of manipulating these microbial communities for human benefit.

Our drive to understand how particular fungi are incorporated or rejected from a microbial community is motivated by the large impact fungal composition can have on the outcome for human and plant health. Fungi in complex polymicrobial infections act as markers of disease severity, particularly in the lungs of patients with cystic fibrosis ^13^. In this environment, *Aspergillus fumigatus* and *Candida albicans* are two opportunistic fungal pathogens that are susceptible to phenazines, yet are routinely isolated from patients who are co-infected with the prolific phenazine producing bacterium *Pseudomonas aeruginosa* ^9,10^. Likewise, fungi play prominent roles in the rhizosphere, where they can help the host plants acquire nutrients and water as well as withstand stress and pathogens ^14,15^; plant-growth promoting fungi such as *Trichoderma* and *Penicillium* species are often found in rhizospheres containing phenazine producing bacteria, yet their growth is inhibited by phenazines ^11,16^. Conversely, the ability of phytopathogenic fungi to enter the rhizosphere is of interest due to fungi being responsible for a third of all lost crops annually ^17^. This is despite phenazines being credited as a primary factor in stopping a variety of fungal phytopathogens from infecting food crops, including pseudomonads that can suppress *Gaeumannomyces graminis* var. *tritici* and *Fusarium oxysporum* f sp. *radicis-lycopersici*, two fungal pathogens of tomato and wheat, respectively ^4,5^. Finally, plant associated fungi known as mycorrhizae play an outsized role in carbon sequestration: mycorrhizae-associated vegetation sequester approximately 350 gigatons of carbon a year compared to 29 gigatons stored by nonmycorrhizae-associated vegetation ^3^. Notably, phenazine producers are found in diverse environments beyond food crops, including in forests and grasslands, thus pointing to another niche of consequence where fungi must navigate phenazine assault ^6^.

How do fungi maintain an active presence in microbial communities where they run the risk of encountering phenazines? Recognizing that some soil bacteria can tolerate phenazines well ^18–20^, we hypothesized that one mechanism by which fungi might gain navigate such hostile environments is through association with a protective bacterial partner. Precedent for such relationships exists. For example, members of the *Burkholderiaceae* family form associations with fungi. *Trichoderma asperellum* is a biocontrol fungus that suppresses the wheat pathogen *Fusarium oxysporum*. *Paraburkholderia terrae* associates with the mycelium of *T. asperellum* and can be induced to migrate in the direction of mycellial growth, as well as promote fungal growth in the presence of crude supernatant derived from antagonistic bacteria ^21^. However, this family of bacteria can also empower pathogenic fungi. *Rhizopus microsporus* is a necrotic plant pathogen of rice. The primary toxin it secretes that is required for infection is actually produced by the intracellular bacterium *Paraburkholderia rhizoxinica* that resides inside the fungal cells ^22^. Other *Paraburkholderia* with less clear roles associate extracellularly with fungal pathogens, such as *P. fungorum*, found isolated with the white-rot fungus *Phanerochaete chrysosporium* ^23^. While the roles each of these bacteria play for their host fungus may differ, fungal association with bacteria of this family is well established.

In addition to these isolated examples, data from a recent metagenomic survey of soil microbes across many climate conditions support the notion that cooperation with bacteria might underpin fungal ecological success ^24^. Specifically, this study found that the presence of bacterially-derived genes regulating antibiotic tolerance were correlated with fungal biomass in the community. Although this co-occurrence was suggested to indicate inter-domain antagonism, where bacterial groups use these genes to defend themselves against fungally-produced antibiotics, an alternative and non-mutually exclusive explanation may be that these bacterial stress response genes help fungi navigate an otherwise inhospitable environment. Given that fungi can be excluded from microbial communities by phenazine-producing bacteria, it stands to reason other phenazine resistant bacteria in the community that associate with fungi may have the power to affirm their presence. Because the number of environments where susceptible fungi are found living in proximity to phenazine producers is likely to be high, we reasoned that finding an example of such a hypothetical protective association could be of great value in understanding the recruitment versus repression of fungi in microbial communities containing phenazine producers.

Accordingly, we set out to identify a model bacterial/fungal partnership in the presence of phenazines. Using an accessible soil on the Caltech campus from which we had previously isolated phenazine degrading bacteria ^19^, we designed a procedure to select for such partnerships. Here, we report the isolation and initial mechanistic characterization of a genetically tractable fungal-bacterial system where the bacterial partner protects the fungus from PCA assault. The ease with which we were able to experimentally validate the existence of a hypothetical bacterial/fungal association suggests that this type of partnership may be widespread in nature.

## Results

### Isolation of protective bacterial partner and physically associated fungus

To identify fungi that resist phenazine assault with protective bacterial partners, we sampled topsoil from the base of a blood orange citrus tree outside of the Beckman Institute on the Caltech campus. We chose this site because soil represents an easily accessible and broad niche containing many microbial species and because we had isolated a strain of *Mycobacterium* from this same plot that can degrade phenazines, suggesting the presence of bacteria capable of producing and interacting with these molecules ^19^.

We collected the top three centimeters of soil from this site, and developed a protocol to find strong bacterial-fungal pairs. We washed and sonicated 100 mg to separate microbes that were not strongly associated with one another, thereby enriching for strongly adherent partners (**Fig. 1A**). To select a first fungal culture, the washed samples were diluted to extinction and plated on potato dextrose agar. Fungal colonies that grew after approximately three days were screened for the presence of bacteria via PCR amplification of the 16S rDNA region. Fungal/bacterial pairings were then challenged with 300 μM phenazine-1-carboxylic acid (PCA). We used PCA because it is the biosynthetic starting product for modification into more specialized phenazine types and is known to play important roles in excluding fungal pathogens from wheat rhizosphere communities ^5^. Co-colonies that were able to grow when challenged with PCA were repeatedly sub-cultured to isolate the partner bacterium, while the fungus was re-plated in the absence of PCA with bacteriocidal antibiotics to cure it of the bacterium. Isolated fungi and bacteria were retreated with PCA to check phenazine-sensitivity and tolerance, respectively, and susceptible fungi were then supplemented with their co-isolated bacterium in the presence of PCA to confirm that the partner bacterium conferred phenazine tolerance.

**Fig. 1.**
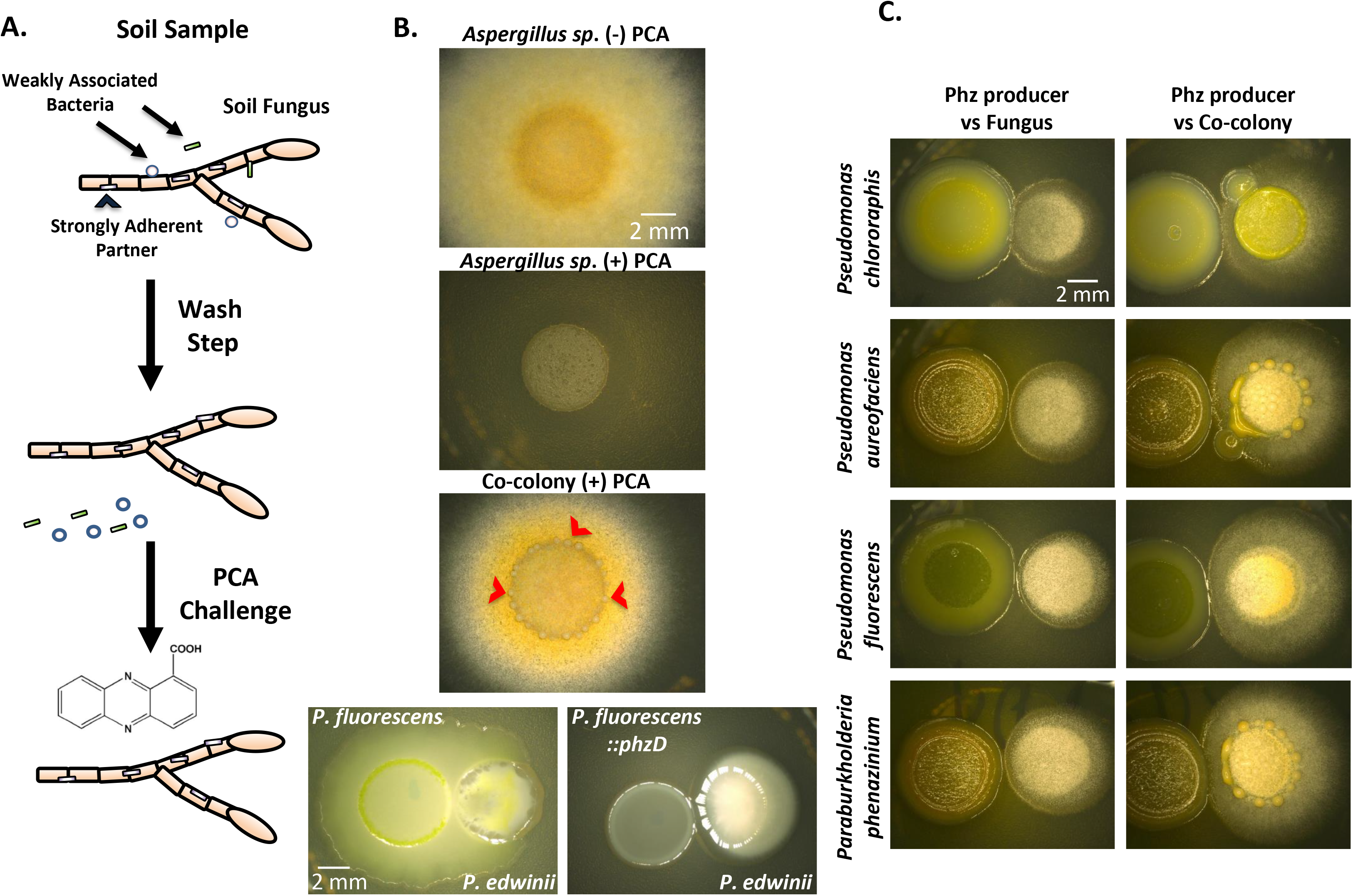
Co-isolation of a fungus and protective bacterial partner. **A)** 100 mg of top soil samples were washed in PBS with 0.01% Tween 20, sonicated to break apart soil and loosely associated microbes, and plated on PDA. Colonies were screened for bacterial partners by 16S amplification, and pairings were subsequently cured of their partners and tested by PCA challenge. **B)** An *Aspergillus* isolate from the soil growing on PDA (column, top). The same isolate fails to grow in the presence of 300 μM PCA after 48 hours (column, center), but is capable of withstanding phenazine assault when grown with its co-isolated partner, *P. edwinii* (column, bottom). When *P. edwinii* is grown alone next to *P. fluorescens*, it can be engulfed by the phenazine-producing strain (bottom row, left) but not the phenazine mutant strain, ::*phzD* (bottom row, right). **C)** The isolated *Aspergillus* species is inhibited by a wide array of phenazine producing organisms (left column), The bacterial competitors and primary phenazines they produce, top to bottom, are *Pseudomonas chlororaphis* (phenazine-1-carboxamide), *Pseudomonas aureofaciens* (2-hydroxyphenazine), *Pseudomonas fluorescens* (PCA), and *Paraburkholderia phenazinium* (iodinin). Fungal growth is enhanced in the same conditions by the presence of *P. edwinii* (right column). Also note *P. fluorescens* is incapable of engulfing the co-colony as it did to *P. edwinii* alone in B. Bacterial aggregates are visible in several images. All colonies were grown on PDA for 48 hrs at 30 °C.

Using this process we uncovered three fungal-bacterial partnerships. Genus-level identification was performed with ITS and 16S rDNA sequencing, respectively. Two *Paraburkholderia* isolates were found protecting an *Aspergillus* and a *Lecythophora* isolate. A *Luteibacter* species also provided protection to a second *Aspergillus* isolate (**Fig. S1A**). In each case, the fungal growth was negatively impacted when challenged with PCA alone, but was restored to varying degrees when supplemented with its natively co-isolated bacterial partner. Of these three co-isolates, we selected the *Paraburkholderia*-*Aspergillus* pairing for further analysis due to the dramatic level of protection the bacterium provided the fungus, as well as the radical morphological change the bacterium underwent as it formed spherical aggregates within the fungal colony when the two were challenged with PCA (**Fig. 1B**). Moreover, while the *Aspergillus* species is sensitive to PCA and *P. edwinii* resists its toxic effects, *P. edwinii* is still vulnerable to engulfment by the PCA producer *P. fluorescens*, but not to a strain that cannot make PCA, suggesting a mutual benefit (**Fig. 1B**). Finally, this pairing was attractive because previous reports of *Burkholderiaceae* family members being isolated with fungi ^21,25^ suggested that such associations may be common in the soil. Due to this bacterium’s ability to help its fungal partner prosper despite the presence of a toxin, and its phylogenetic placement, we named it *Paraburkholderia edwinii*, derived from the Old English “Edwin”, meaning prosperous friend.

### *P. edwinii* protects its fungal partner from phenazine assault

We next sought to characterize the range of the bacterium’s ability to protect its partner fungus from phenazines. The minimal inhibitory concentration of PCA toward fungal targets has been reported to be in the 1-50 μM range ^26^. The 300 μM PCA used for our isolation assay therefore represents a strong phenazine challenge intended to identify bacterial partners with a robust protection phenotype. The advantage of using a high concentration of PCA in our laboratory experiments is that it may better mimic local gradients of PCA that exist within rhizosphere microbial communities that likely exceed bulk measurements.

To determine whether *P. edwinii* can protect *Aspergillus* against actual phenazine-producers in addition to purified PCA, we tested the response in the presence of different phenazine-producing *Pseudomonads*. *Aspergillus* was plated adjacent to *Pseudomonas fluorescens*, *Pseudomonas chlororaphis*, *Pseudomonas chlororaphis* sub-species *aureofaciens*, and *Paraburkholderia phenazinium*. The primary phenazine product made by these species under are growth conditions is PCA, phenazine-1-carboxamide (PCN), 2-hydroxy-phenazine, and iodinin, respectively. When grown close to one another, each phenazine producing bacterium impeded *Aspergillus* growth (**Fig. 1C**). However, when the fungus was supplemented with *P. edwinii*, growth was partially restored. Intriguingly, bacterial aggregates again formed within the co-colonies proximal to phenazine producers, suggesting this morphological phenotype reflected a general protective response (**Fig. 1C**). Finally, to verify that these responses were specifically due to phenazine assault, mutants of *Pseudomonas fluorescens* and *Pseudomonas chlororaphis* were obtained that could not make phenazines ^27^. While the WT strains were capable of suppressing fungal growth, the fungus grew unimpeded in proximity of the non-phenazine producing mutants (**Fig. S1B**), confirming that the protection provided by *P. edwinii* is specific to phenazine assault and can occur in a mixed microbial system.

### *P. edwinii* undergoes a morphological shift in response to phenazine-induced fungal stress

To understand how *P. edwinii* responds to its partner fungus during phenazine assault, we imaged the bacterium inside the co-colony. While a ring of what appeared to be one or two dozen bacterial aggregates formed on the co-colony surface, it remained possible these were fungal structures. To distinguish between these possibilities, we adapted a tissue clearing technique developed in our lab termed Microbial identification after PASSIVE Clarity Technique (MiPACT) to render the fungal tissue transparent (see Materials and Methods). This allowed us to visualize bacteria within the fungal structure using *in situ* fluorescence detection of 16S rRNA with the hybridization chain reaction (HCR) ^28^. Because the exterior of the colonies showed putative bacterial aggregates within the center, we hypothesized that may be where the bacteria were concentrated. We first imaged the outer 2/3 of the colonies. In the control, whole-colony samples untreated with PCA, *P. edwinii* was found concentrating near the tips of the outwardly-growing mycelium, with relatively low amounts of bacteria found further inward among older mycelial growth. The propensity of *P. edwinii* to track with the mycelial edge in the untreated samples is in agreement with reports suggesting other *Paraburkholderia* are capable of identifying the growing edge of expanding fungi ^21^. Conversely, in the PCA treated sample, this population of bacteria was largely absent (**Fig. 2**).

**Fig. 2.**
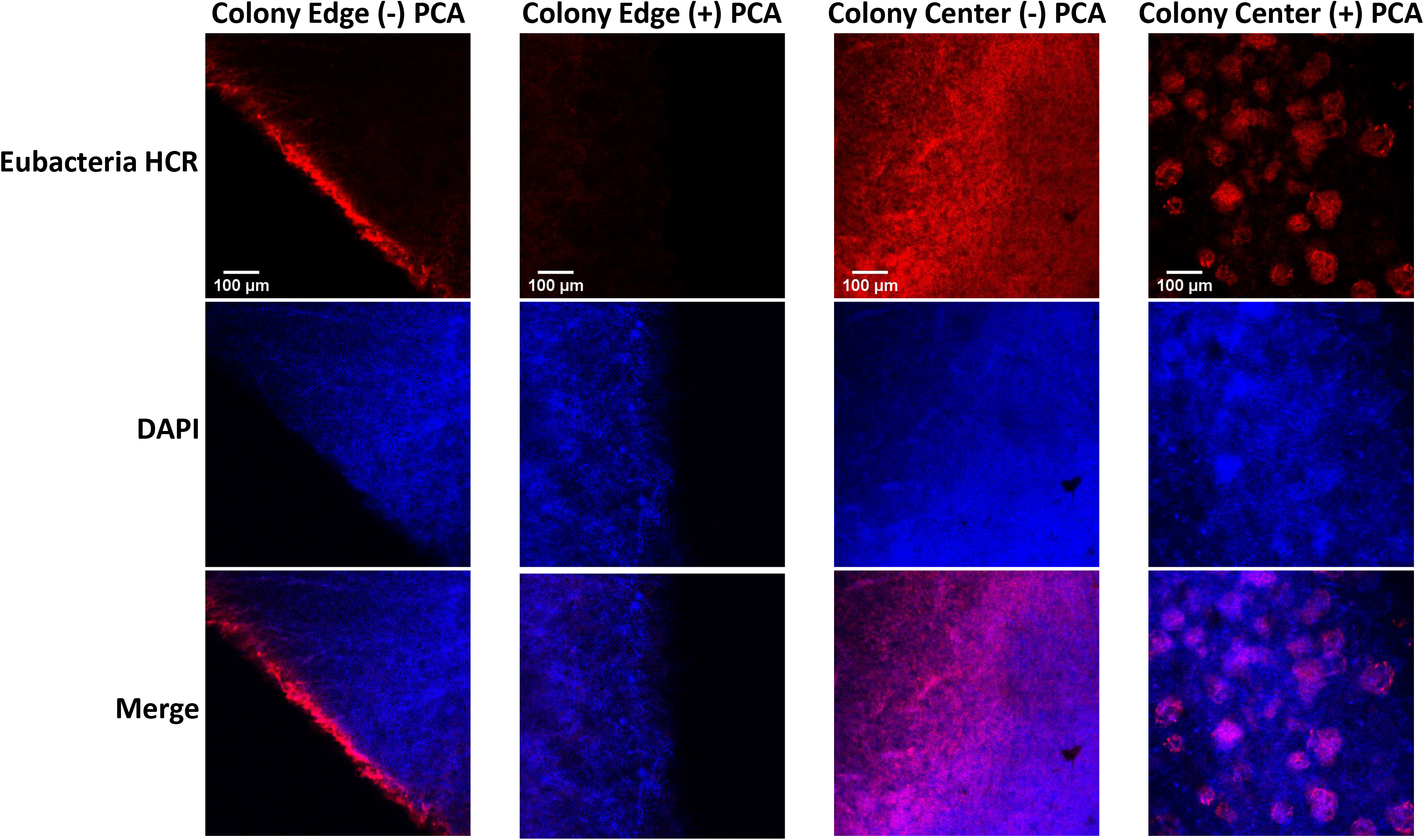
*P. edwinii* forms aggregates in center of fungal colony in response to PCA challenge. Bacteria gather along the edge of the mycelia (left column) and mixed throughout the interior of the fungal colony (right center column). When challenged with PCA, *P. edwinii* forms congregates less at the leading edge of the mycelium (left center column) but aggregates within the colony center (right column). Whole colonies were grown on PDA for 48 hours at 30 °C, processed using the MiPACT technique to render fungal tissue transparent, and visualized using HCR eubacterial probes and DAPI (see materials and methods). Eubacterial probes were labeled with a with an Alexa 647 fluorophore. Images are representative of three independently grown co-colonies for each condition, and were captured on an inverted confocal Leica model TCS SPE confocal microscope with a 10x objective. Images of the HCR signal were normalized in contrast to the brightest image in like samples (i.e. edge vs edge, or center vs center), while images of the DAPI signal were independently adjusted to best outline fungal morphology in the vicinity of the bacteria.

To gage whether a lack of bacteria among the outer mycelium in the PCA treated samples was due to bacterial aggregation, we imaged the center of each co-colony. In untreated samples, bacteria were mixed homogenously throughout the fungal mass without identifiable structure or patterning **(Fig. 2**). One exception to this observation was an apparent transition zone between the bacterially rich inner co-colony and more sparely populated outer co-colony. This transition zone, or ring, comprised more densely packed bacteria, but the region lacked further organization (**Fig. S2**). In the PCA treated sample, the center of the co-colony contained clear spherical structures that lit up with the eubacterial HCR probes (**Fig. 2**). These *P. edwinii* aggregates ranged from 50 to 100 μm in diameter and were ubiquitous throughout the colony center, indicating that the visible bacterial aggregates on the surface of the co-colony were only a fraction of those being formed. The PCA treated co-colony also had a transition ring structure between the bacterially populated center and unpopulated outer colony. In this case, however, the ring contained more clearly defined aggregates of bacteria, corresponding to the region that produced aggregates visible to the naked eye (**Fig. S2**). These results reveal that *P. edwinii* forms bacterial aggregates inside and in the center of the fungal colony when challenged with PCA.

### *P. edwinii* acts as a toxin sponge

How does *P. edwinii* offer resistance from phenazine assault to its partner fungus? Possible mechanisms included phenazine degradation, sequestration, and/or detoxification. We first tested degradation. To accurately measure the PCA concentration over time, we grew *P. edwinii* in shaking liquid cultures spiked with 300 μM PCA either alone or in the presence of the *Aspergillus* species, in case a fungal signal was necessary to trigger degradation of PCA. In no condition was PCA degraded (**Fig. S3**). The lack of degradation suggested that bacterial aggregate formation might instead reflect a PCA sequestration and detoxification response, which we proceeded to test.

Because the bacterial aggregates are too small to probe or manipulate individually, we modified our experimental set up to grow *P. edwinii* directly next to its *Aspergillus* partner in the presence of PCA to generate bacterial auto-aggregation in the form of a colony. To verify the protection phenotype is still responsive in this assay, we grew the two organisms next to each other in the presence and absence of PCA. Growing *P. edwinii* and the *Aspergillus* species at a distance in the presence of PCA resulted in severely stunted fungal growth, however the fungus was able to grow toward *P. edwinii* when plated adjacently (**Fig. 3A**). Intriguingly, the bacterial colony developed a deep yellow hue in the PCA treated condition, but only did so in the presence of the fungus. PCA is a largely colorless molecule when exposed to oxygen, but in the reduced state turns yellow. We used LC-MS to determine whether this yellow pigment was PCA and its presence was confirmed in the bacterial sample grown next to the fungus (**Fig. 3B**). We detected a smaller amount of PCA in the colonies of *P. edwinii* grown alone in the presence of PCA than in the presence of PCA and the fungus, suggesting that while PCA sequestration may be an intrinsic trait of the bacterium, sequestration is stimulated by the fungal partner (**Fig. 3C**).

**Fig. 3.**
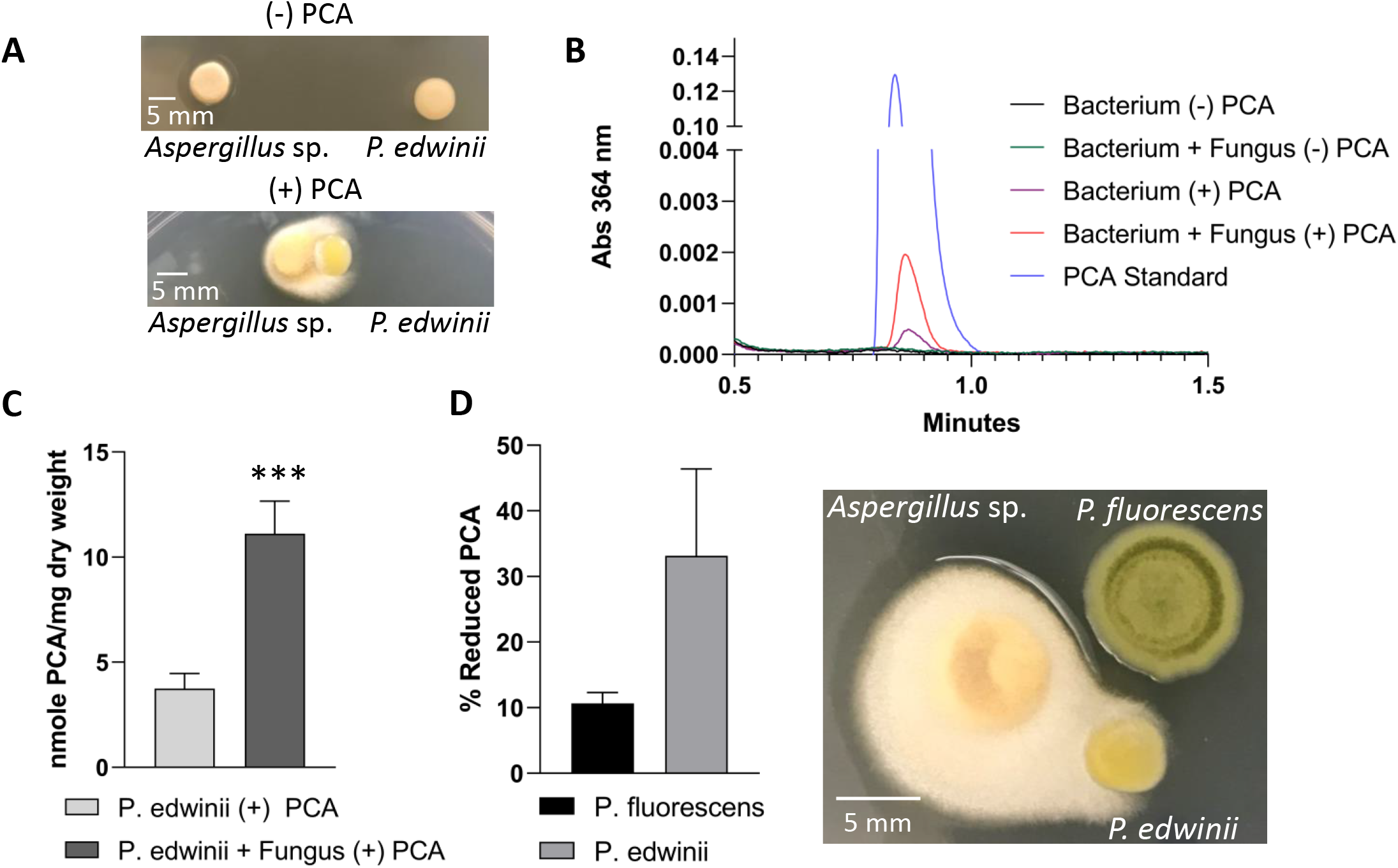
*P. edwinii* acts as a toxin sponge. **A)** The *Aspergillus* species and *P. edwinii* growing on PDA supplemented with 300 μM PCA growing at a distance (top) and 5 mm apart (bottom). Note the *Aspergillus* growth toward the bacterium and deepening yellow color of *P. edwinii* when the two organisms are grown in proximity of one another. Images are representative of 5 sets of colonies. **B)** HPLC chromatogram with the peak representing PCA derived from scraped up *P. edwinii* colonies grown with or without its fungal partner, and in the presence of absence of PCA challenge. **C)** nmoles PCA sequestered by *P. edwinii* in the absence and presence of its partner fungus, normalized by bacterial dry mass. Quantification was performed by measuring absorbance at 365 nm. Error bars represent standard deviation of four biological replicates. *** p < 0.001. **D)** Fraction of reduced PCA present (left) in a *P. edwinii* or *Pseudomonas fluorescens* colony when grown as a three-member system with the isolated *Aspergillus* species (right). Reduced PCA was quantified by fluorescence spectroscopy using an excitation wavelength of 365 nm and reading emission at 520 nm. Error bars represent standard deviation of 3 biological replicates.

Having confirmed the presence of PCA within the bacterial colonies, we next wanted to assess its redox state. Previously, we had been measuring PCA sequestered from an agar plate where most of the molecule would be expected to be oxidized due to atmospheric oxygen. A better comparator for the fraction of reduced PCA found within a *P. edwinii* colony would therefore be another bacterial colony containing PCA, but one that was not employing a protection response involving PCA detoxification. To this end, we grew *P. edwinii* and the *Aspergillus* species adjacent to the phenazine producer *P. fluorescens*. PCA is the sole phenazine produced *by P. fluorescens*. Before the bacterial colonies were harvested, the plates were transferred to an anaerobic chamber to minimize the atmospheric oxidation of any reduced PCA. Reduced, but not oxidized, PCA has a peak excitation of 364 nm and emission of 520 nm, which allowed us to determine the fraction of the PCA in the reduced state. Fluorescent emission spectra were collected in the anaerobic chamber, which revealed the phenazine-producing *P. fluorescens* colony biofilm to contain approximately 10% of its PCA in the reduced state whereas *P. edwinii* colonies maintained approximately a third of the PCA it sequestered in the reduced state. This result demonstrated that *P. edwinii* colony aggregates reduce PCA (**Fig. 3D**).

### HrcA is a regulator of the protection response in *P. edwinii*

Given that PCA sequestration in *P. edwinii* is stimulated by its partner fungus, we aimed to discover how *P. edwinii* sensed and responded to its partner. We developed genetic tools to manipulate *P. edwinii* to screen for mutants altered in their ability to protect the fungus from PCA. Because *P. edwinii* is a novel soil isolate, we sequenced its genome using Illumina and PACBio technologies. Two closed chromosomes resulted upon assembly. Because most members of the *Burkholderiaceae* family contain an additional third genetic element that can range in size from one or more plasmids to a small third chromosome, we attempted to also isolate smaller genetic components. However, unlike many closely related species of *Paraburkholderia*, no plasmid was recovered. *P. edwinii* appeared to be a novel species, with the closest match being *Paraburkholderia SOS3*, with 88% average nucleotide identity shared between the two. We also sequenced the *Aspergillus* species, and we report an assembly of 26 contigs greater than 0.5 MB accounting for ~38 MB. See materials and methods section for more details.

We developed a mating protocol to introduce a mini-mariner transposon ^29^ into the genome of *P. edwinii*. Mutants were screened with the *Aspergillus* species on PCA, and mutants that produced an atypical morphology had their transposons mapped to the inserted gene (Supplementary Table 1). *P. edwinii* mutants were identified that were either less or more protective of the partner fungus. Generally, more bacterial aggregation was associated with more fungal protection and vice versa (**Fig. 4A**). We focused on a mutant with a transposon insertion in the stress inducible transcriptional repressor *hrcA*, due to its strong protection phenotype and relative ease of growth. To ensure its phenotype was due to the disruption of this gene and was not caused by polar effects or secondary mutations elsewhere in the genome, we made an in-frame deletion of *hrcA*. The deletion mutant phenocopied the transposon mutant, showing enhanced bacterial aggregation and fungal growth when challenged with PCA (**Fig. 4B, C**). Constitutively expressing *hrcA* from a pBBR1 vector restored the WT phenotype **(Fig. 5B**). Not only did the Δ*hrcA* mutant promote fungal growth beyond the wild type strain during PCA challenge, but the extra-large bacterial aggregates characterizing this mutant appeared to form even in the absence of PCA, suggesting this mutant constitutively turned on the protection program (**Fig. S4A**).

**Fig. 4.**
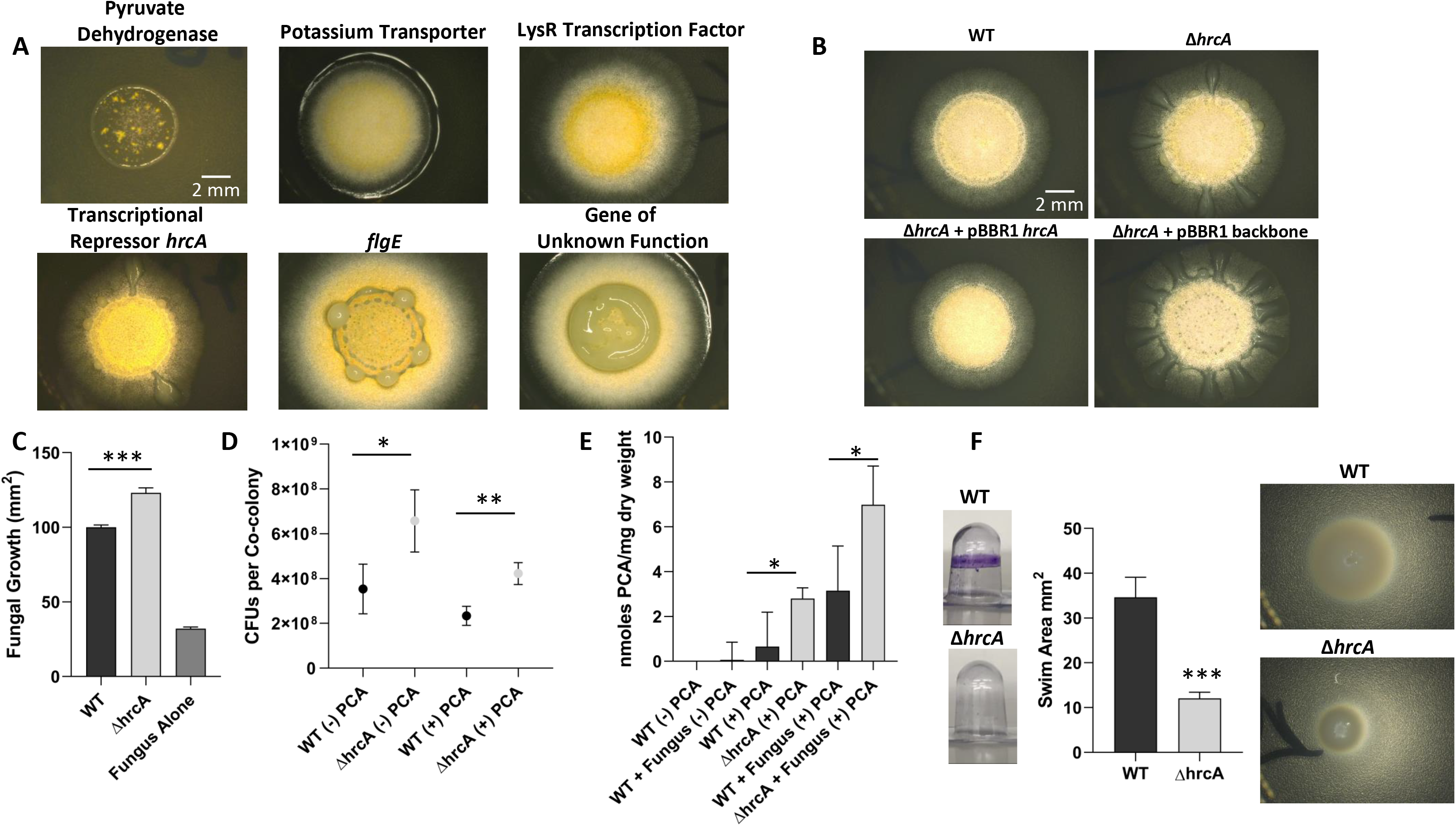
*hrcA* regulates the protection response of P. edwinii. **A)** Transposon mutants of *P. edwinii* that alter the fungal protection response when grown for 48 hours on PDA supplemented with 300 μM of PCA, organized from less bacterial aggregation/protection to most. **B)** An in-frame deletion of the *hrcA* gene was constructed to verify the transposon phenotype, and was complemented using the pBBR1 expression vector. **C)**Δ*hrcA* shows a greater degree of fungal protection compared to the WT strain. Error bars represent standard deviation of four biological replicates. ***p < 0.001. **D)** Comparison of WT and Δ*hrcA* CFUs derived from co-colonies. Whole co-colonies exposed or not exposed to PCA challenge were excised from PDA plates, homogenized, and plated on PDA supplemented with nystatin to prevent fungal growth. Reported is the total number of CFUs per co-colony. Error bars represent standard deviation of four biological replicates. *p < 0.05, **p < 0.01. **E)** Comparison of the ability of WT and Δ*hrcA* to sequester PCA with and without partner fungus. Error bars represent standard deviation of four biological replicates. *p < 0.05. **F)** Biofilm and motility assay of WT and Δ*hrcA*. The WT strain shows increased biofilm formation and motility (top images) compared to Δ*hrcA* (bottom images). The biofilm formation assay utilized 1/5 V8 medium and was grown for 24 hours at 30 °C before staining with 0.1% crystal violet, while the motility assay was conducted in modified M9 medium for 72 hours at 30 °C. Error bars represent standard deviation of four biological replicates. ***p < 0.001.

**Fig. 5.**
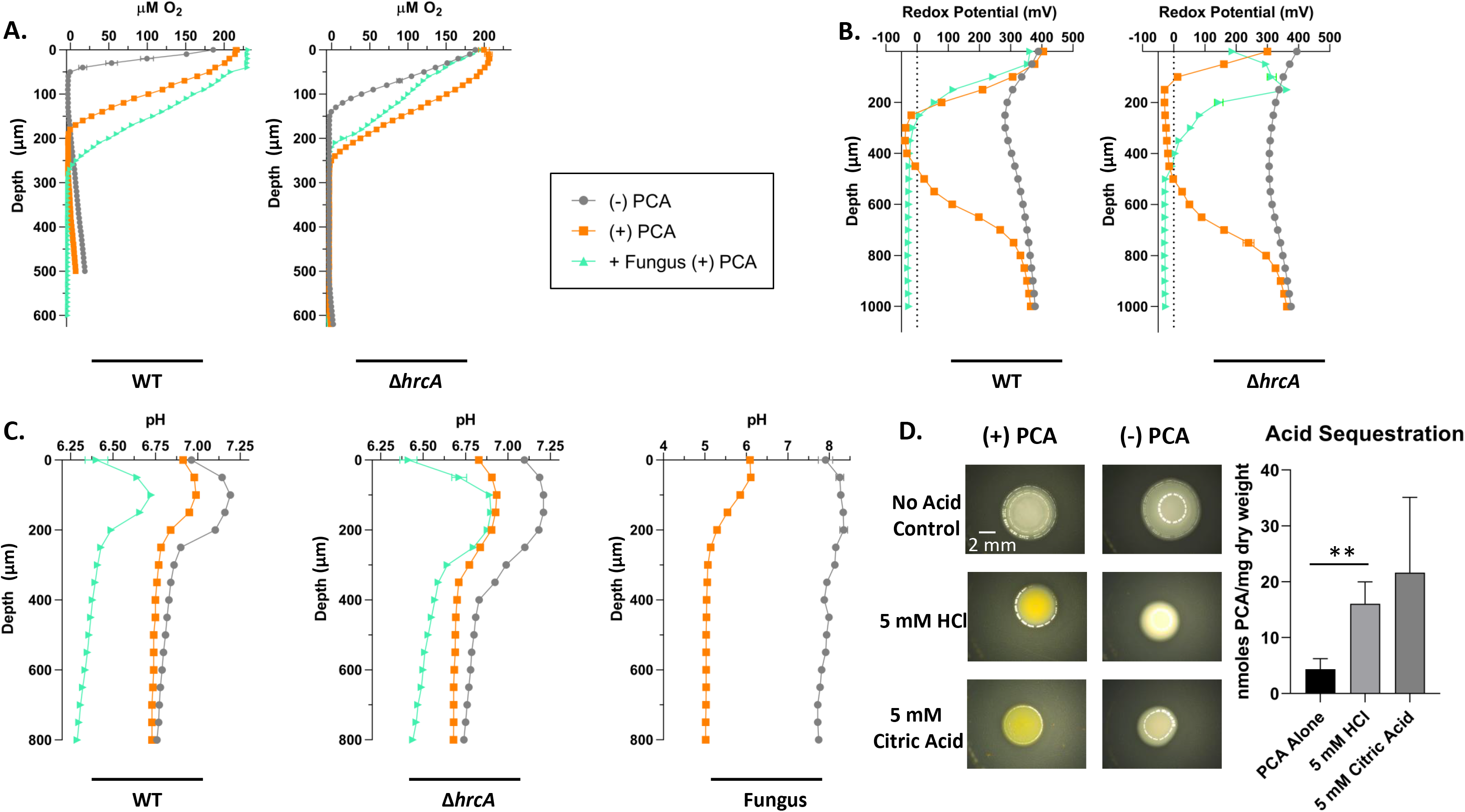
*P. edwinii* and *Aspergillus* species respond to PCA challenge by modifying oxygen availability, reduction potential, and pH. **A)** Oxygen profile of *P. edwinii* colonies. The WT colony grows as a flat disc in the absence of PCA, but becomes rounded with an apparent outer polysaccharide layer when stressed with PCA, and the increase in colony volume creates a larger anoxic zone beneath this layer than exists in the non-stress condition (left). Δ*hrcA* shows a similar trend but more closely resembles the WT PCA (+) condition even when not challenged (right). **B)** When challenged with PCA, WT and Δ*hrcA P. edwinii* colonies generate more reducing conditions. The larger zone of lower reducing potential in the Δ*hrcA* mutant compared to WT when exposed to PCA is reflective of the larger internal volume this mutant creates. The addition of the *Aspergillus* isolate causes a further decrease in reduction potential that is sustained to greater depths (left right)**C)** The WT and Δ*hrcA P. edwinii* colonies generate near-neutral pH conditions that show a trend of decreasing when exposed to PCA, and growing the fungus adjacent to *P. edwinii* colonies causes the internal environment to be more acidic (Left, Center). The *Aspergillus* isolate generates alkaline conditions when grown without PCA challenge, but generates a much more acidic environment when PCA is present (Right. Error bars for all microelectrode experiments represent standard deviation of three measurements at each depth. **D)** Addition of 5 mM HCl or citric acid causes an increase in PCA sequestration in *P. edwinii* colonies. Colonies were grown for 48 hrs at 30 °C. All adjacent colonies were grown 5 mm apart. Error bars represent standard deviation of four biological replicates. **p < 0.01.

Intriguingly, the *hrcA* gene product in *Bacillus subtilis* represses expression of genes involved in the stress response to heat shock, and deletion of *hrcA* in *B. subtilis* results in cells that can adapt and grow more rapidly under conditions of heat stress ^30^. By analogy, we wondered whether the removal of *hrcA* in *P. edwinii* might permit larger aggregate formation due to a similar growth advantage in the presence of PCA. To test this, we grew WT and Δ*hrcA* strains as co-colonies with the *Aspergillus* species in the presence or absence of PCA. Co-colonies were homogenized after 48 hours and bacterial CFUs were plated on potato dextrose agar containing nystatin to suppress fungal growth. The Δ*hcrA* strain showed an approximately 2-fold increase in CFUs compared to the WT per co-colony in both the PCA treated and untreated conditions (**Fig. 4D**). Not only did the Δ*hcrA* mutant grow better than the WT in the presence of PCA, it was better at sequestering PCA; as before, its ability to sequester PCA was stimulated in the presence of the fungus (**Fig. 4E**). Because the amount of PCA sequestered is normalized by the dry weight of the collected biomass, a growth advantage alone is not sufficient to explain these results. Though bacterial aggregation in the Δ*hcrA* mutant was enhanced relative to the WT, it failed to form a biofilm using the crystal violet assay ^31^—possibly linked to a swimming motility defect we uncovered using a swim assay(**Fig. 4F**). These results suggest that the protection/aggregation phenotype relies on a different developmental program from that involved in classical biofilm development.

### *P. edwinii* holds sequestered PCA in a reduced, anoxic environment

Phenazines exert toxic effects on diverse cell types through a variety of mechanisms including generating reactive oxygen species and destabilizing the electron transport chain of the target cell dependent upon the ability of the phenazine to cycle its redox state in the presence of oxygen ^13,32–37^. It may therefore seem paradoxical that the *Aspergillus* species achieves protection against PCA by promoting concentration of PCA within its co-culture. Given that *P. edwinii* colonies are enriched in reduced PCA relative to the phenazine producer, *P. fluorescens* (**Fig. 3D**), we hypothesized that a solution to this paradox might come from limiting oxygen by maintaining a reducing environment within the bacterial aggregates. We used oxygen and redox microelectrodes to test this hypothesis.

Because the co-colony bacterial aggregates are of a similar size as the width of our microelectrode tips (25 – 100 μm), puncture when probed, and are invisible from the outside of the co-colony, we instead grew *P. edwinii* next to the *Aspergillus* species with or without PCA to induce the sequestration phenotype and probed the interior of the bacterial colony (setup as in Fig. 4A). We measured the oxygen concentration and redox potential of the bacterial colony microenvironment after 48 hours (**Fig. 5**). In the absence of PCA, *P. edwinii* grew as shallow, flat colonies with a narrow anoxic zone Fig. S4B). These colonies were anoxic at 50 μm of depth before increasing in oxygen concentration at approximately 175 μm depth (**Fig. 5A**). When grown without the fungal partner in the presence of PCA, the bacterial colony became taller and encased in an apparent layer of polysaccharide (**Fig. S4B**). Oxygen levels in this thicker colony dropped slowly through the outer layer, disappearing at approximately 180 μm. Oxygen was again detected at 380 μm, indicating a larger anoxic volume within the colony compared to the untreated colony. When grown next to its partner fungus in the presence of PCA, these trends continued with the colony again becoming taller and more dome shaped (**Fig. S4B**), becoming anoxic at 280 μm beneath its thicker layer of matrix. At no depth probed was oxygen again detected.

Given the additional protection from PCA the Δ*hrcA* mutant provides, we speculated that it might contain a larger anoxic core even in the absence of PCA challenge/its fungal partner. When grown alone in the absence of PCA, Δ*hrcA* grew tall, rounded colonies (**Fig. S4B**) and reached anoxia 150 μm from the surface and continued to a depth of 590 μm (**Fig. 5A**). In the presence of PCA, anoxia was reached at a depth of 250 μm and continued to 660 μm, which resulted in similarly large anoxic interiors even as PCA caused an increase in matrix material at the surface of the colony (**Fig. S4B**). Intriguingly, the presence of the fungus and treatment with PCA resulted in the Δ*hrcA* mutant reaching anoxia at a similar depth as the PCA treatment alone at 230 μm, but the oxygen concentration declined more sharply. Oxygen again could not be detected at any depth when the fungus was present (**Fig. 5A**).

Profiles using a redox probe revealed that, in the absence of PCA, neither WT nor Δ*hrcA* significantly lowered the redox potential through their depth, maintaining a potential of greater than 280 mV in both cases, indicating an oxidizing environment. (**Fig. 5B**). This is not surprising because the absence of oxygen is necessary but not sufficient to create a reducing environment. When supplemented with PCA (the effective redox buffer), however, both strains showed a marked drop in redox potential to a low of approximately −30 mV, although the Δ*hrcA* mutant maintained low redox potential over a greater depth and thus represents a larger volume of a reducing environment. When challenged with PCA in the presence of the fungus, both strains showed similar low redox potentials in their cores as when challenged with PCA alone, and the environment continued to be reducing at all tested depths for both strains. To determine if the *Aspergillus* species contributes to this reducing environment, the redox potential of fungal colonies was measured with and without PCA challenge. The fungus showed a sharp drop in redox potential when challenged with PCA (**Fig. S4C**). Although the drop was somewhat less than that produced by the bacterium, it is possible that the fungus helps maintain a reducing environment in partnership with *P. edwinii*.

### Fungal-induced pH shift corresponds with protection response

While the *Aspergillus* species stimulated *P. edwinii* to generate anoxic and reducing interiors in the presence of PCA, the fungus was not required to trigger these bacterial responses. However, the *Aspergillus* species promoted an increase in PCA sequestration in both WT *P. edwinii* and even more so in the Δ*hrcA* strain (**Fig. 4E**), suggesting the fungus may provide a stress-related trigger to the bacterium. Many species of fungi will acidify their environment when stressed in an attempt to outcompete other microbes ^38^. Accordingly, we hypothesized that our *Aspergillus* species might acidify the medium in response to PCA. In addition to the acid stress to which this would expose the bacterium, a lower pH results in a higher fraction of the PCA becoming protonated, and thus neutrally charged and more cell permeable, potentially forcing a response from *P. edwinii*. To test this hypothesis, we used a pH microelectrode to probe the pH of *P. edwinii* and *Aspergillus* colonies

Both the WT and the Δ*hrcA* strain exhibited a colony pH profile above 7.0 when grown alone without PCA, with a slightly more acidic pH profile in the presence of PCA and an even more acidic profile in the presence of PCA + the fungus (**Fig. 5C**). Measurement of pH in fungal colonies alone showed that PCA exposure prompts the fungus to dramatically acidify its environment by 2-3 log units (**Fig. 5C**). Could acidification be a trigger for the protective response of *P. edwinii*? Given the dual stress induced by acidification in the presence of PCA and the involvement of a stress regulator in the activation of the protective PCA sequestration and reduction phenotype, we hypothesized that acidifying the medium could cause *P. edwinii* to behave as though its partner fungus is present even when absent. Indeed, HCl-acidified medium caused *P. edwinii* alone to sequester nearly four-fold more PCA, an action that previously required the fungal partner (**Fig. 5D**). We obtained similar results with citric acid, an organic acid made by some *Aspergillus* species and also commonly excreted from roots in the rhizosphere.

### Bacterial Protection of Fungi is not Specific to a Single Species Pairing

To determine if mechanisms underpinning the protective partnership are general enough to allow other *Paraburkholderia* species to protect the *Aspergillus* isolate, we assayed the protective phenotype of three other *Paraburkholderia* species: *P. SOS3*, *P. unamae*, and *P. phenazinium*. *P. SOS3* was isolated in Australia and is genetically similar to *P. edwinii*; *P. unamae* was isolated from the corn roots in Mexico, making it a bonafide member of a food crop rhizosphere ^39^; and *P. phenazinium*, though inhibitory to our fungal isolate when grown in tandem, is another known rhizosphere member and has the ability to form nitrogen fixing nodules ^40,41^. When challenged with 300 μM PCA, both *P. SOS3* and *P. unamae* protected the *Aspergillus* isolate similar to *P. edwinii*, whereas *P. phenazinium* did not (**Fig. 6A**). If a pH shift helps prime the protective response, we similarly wondered whether *P. edwinii* may also protect other fungi, given that acidification is a general trait of filamentous fungi ^42,43^. Accordingly, we tested the ability of *P. edwinii* to protect fungi from different niches, including a species of *Fusarium* isolated from infected corn seedling as well as clinical samples of the human opportunistic pathogens *Aspergillus fumigatus* and two *Penicillium* species isolated from the lungs of cystic fibrosis patients. While *A. fumigatus* is primarily thought of as a pathogen, many *Penicillium* species are thought to enhance plant growth while also being opportunistic humans pathogens. *P. edwinii* protected *Aspergillus fumigatus*, the *Fusarium* isolate, and one of the *Penicillium* isolates (**Fig. 6B**).

**Fig. 6.**
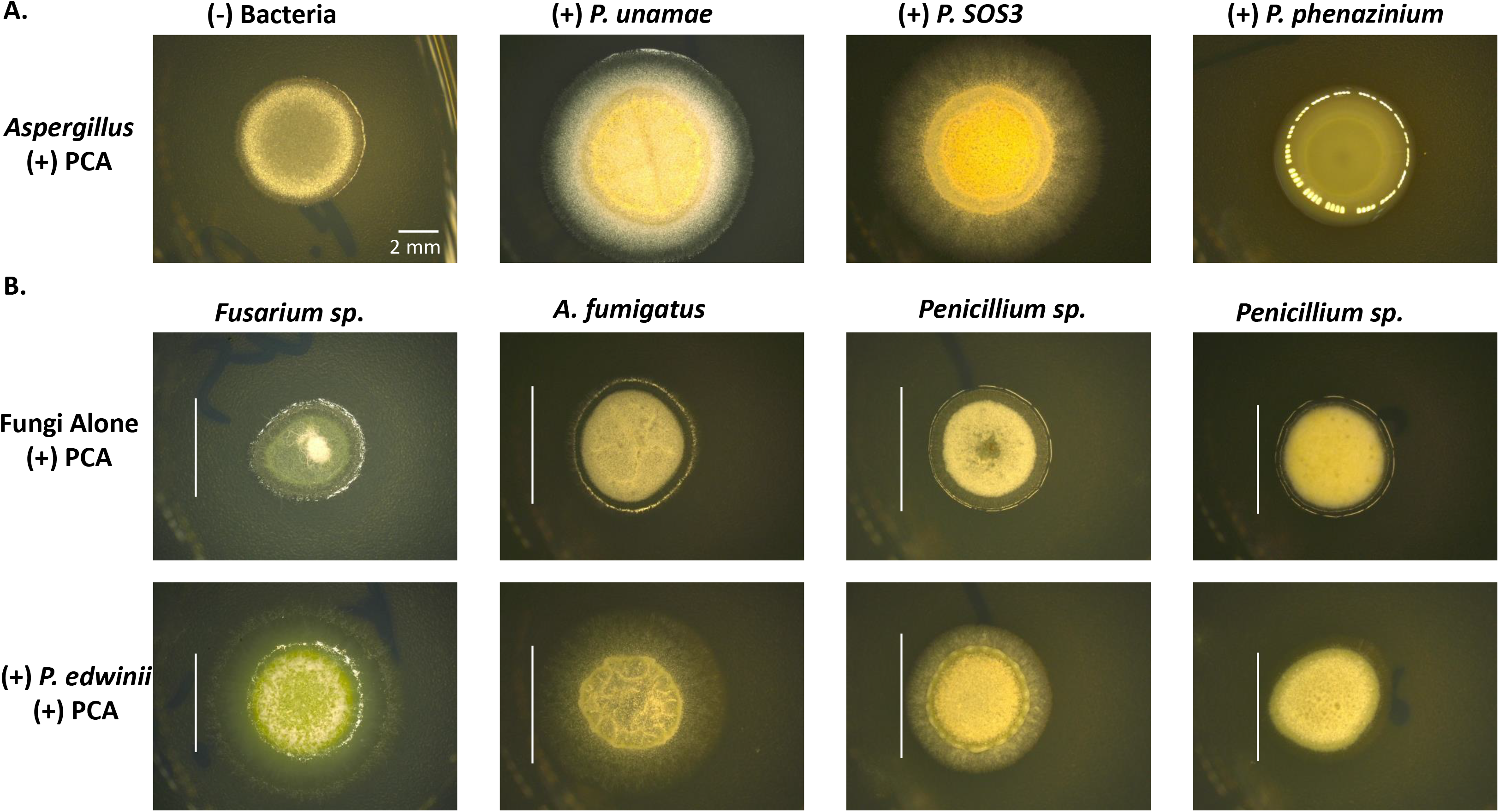
The protection response is conserved in other Paraburkholderia species and shows partial specificity. **A)** The ability to protect the *Aspergillus* isolate was tested among several species of Paraburkholderia. Left to right: No bacterium added, *P. unamae*, *P. SOS3*, and *P. phenazinium*. The protection response was present in *P. unamae* and *P. SOS3*, isolated from the roots of corn in Mexico and from top soil in Australia, respectively. *P. phenazinium*, itself a phenazine producer capable of producing iodinin, appeared to demonstrate no fungal growth. **B)** *P. edwinii* tested for gross ability to protect plant and human pathogenic fungi. From left to right: a phytopathogenic *Fusarium* species isolated in our lab, three opportunistic human pathogenic fungi isolated from the lungs of CF patients including *Aspergillus fumigatus* and two *Penicillium* species. White bars demonstrate the diameter of each fungus in PCA treated condition for comparison with the mixed co-colonies. All samples were grown for 48 hrs at 30 °C.

Together, these results suggest that the ability of bacteria to protect fungi from phenazine assault is not unique to *P. edwinii*. Moreover, the factors involved in activating the protection program are general enough to enable a diverse array of members of this genus to protect a fungus they may not necessarily be paired with in nature, at least under laboratory conditions. The apparent inclusivity of *P. edwinii* between more distantly related fungi but selectivity between members of the same genus (*e.g. Penicillium*) suggests that while the mechanism of protection may be general, there are more factors involved that may help determine the success of such pairings in nature.

## Discussion

This study was motivated by an ecological paradox: how do vulnerable fungi withstand the toxicity of a widespread class of antibiotics (phenazines) produced by co-occurring bacteria in the soil? Given that soil microbial communities are diverse, we hypothesized that other bacteria in these niches would confer protection through inter-domain partnerships. Our discovery of *P. edwinii*—the “prosperous friend” that helps its fungal partner withstand PCA challenge—establishes that such beneficial partnerships exist in nature and are likely common, a finding of basic interest that may also have important practical implications.

While much remains to be learned about the mechanisms underpinning the *P. edwinii-Aspergillus* partnership, our results underscore the importance of the biologically-controlled microenvironment and the biochemical conversation that generates it. *P. edwinii* effectively serves as a “toxin sponge”, sequestering PCA in a reducing environment in response to acidification driven by an *Aspergillus* species. The *P. edwinni* transcriptional repressor HrcA responds to the fungus, triggering the aggregation and PCA sequestration pathways. Yet we do not know how PCA is sequestered by *P. edwinii*—whether it is primarily stored extracellularly in the core of aggregates, or whether some fraction is held intracellularly. Similarly, whether specific enzymes are required to generate and maintain the reducing environment is unclear. A hint at an answer may be found in our mutagenic screen (Table S1): a transposon knockout of the E1 subunit of the pyruvate dehydrogenase gene generated a mutant that actively harms the fungus in a PCA dependent manner, with PCA crystals accumulating in the co-colony (**Fig. 4A**). Intriguingly, pyruvate dehydrogenase has previously been implicated in PCA reduction in *Pseudomonas aeruginosa* ^44^. Future experiments will reveal how the absence of this enzyme promotes PCA toxicity, and whether WT *P. edwinnii* can protect *Aspergillus* against redox-active small molecules other than PCA. That *P. edwinii* is genetically tractable and *Aspergillus* has the potential to be, makes this a good model system to explore these and other mechanistic questions.

Our focus in this study on the Δ*hrcA* mutant derives from the fact that HcrA homologues are stress-inducible transcriptional repressors. For example, heat appears to inactivate HrcA in *B. subtilis*, releasing transcriptional repression of genes responsible for stress-tolerance ^30^. An interesting possibility is that HcrA in *P. edwinii* regulates some of the genes involved in reducing and sequestering PCA in response to its fungal partner. If so, we speculate that HrcA degradation is not due to heat *per se*, but to redox stress from PCA as well as acid or other stress produced by the fungus, where the latter stress could increase the former. We also note that the Δ*hrcA* mutant appears to have its protection program partially “activated” in the absence of these stressors, as the mutant will produce bacterial aggregates within the fungus even without PCA present and will sequester more PCA than WT despite being apart from its fungal partner. That the Δ*hrcA* mutant can sequester still more PCA with the fungus present suggests further regulators or triggers of the protection response await discovery.

An important question raised by our study is whether the phenomenon we have uncovered is environmentally relevant? We believe the answer is yes for several reasons. First, phenazine producers are widespread in nature ^6^ and thus odds are high that fungi will encounter them. Second, we were able to readily isolate a variety of such partnerships. Third, PCA challenge leads to fungal acidification. pH is well known to play a critical role in determining phenazine toxicity as PCA becomes neutrally charged when protonated (pKa = 4.24), leading it to more readily pass through cell membranes ^36,45^. One study investigating the toxicity of PCA found that at a pH of 6.0, PCA showed virtually no toxicity to *C. elegans*, while at pH 5.0 toxicity was very high ^36^. Therefore, while fungal acidification can kill competing microbes, it can also render natural antibiotics made by certain bacteria more toxic. We thus predict that a fungus cooperating with an acid-tolerant beneficial bacterial partner would have a fitness advantage in phenazine-replete microbial communities. Fourth and finally, members of the *Burkholderiaceae* family are known to be particularly acid tolerant, which underlies their ability to associate with fungi ^46^. This, along with phenazine resistance among several individual species, make these bacteria ideal partners to provide protection from phenazine assault to organisms which produce acid in response to stress. Intriguingly, many plant roots also produce organic acid exudates that may reinforce such partnerships. Identifying which organic acids along with other stress signals protective bacteria sense and respond to will be necessary for better understanding and predicting the environmental relevance of these type of bacterial-fungal partnerships.

Given that PCA toxicity and production is predicted to increase in acidic soils that are vulnerable to climate-induced drought due to enhanced oxygen penetration ^8,47^, finding biological agents that can protect fungi from phenazine toxicity may be relevant to agriculture. Many phenazine-sensitive filamentous fungi in the rhizosphere play important roles in water and nutrient acquisition for their host plants, and can help them withstand environmental stresses. Many *Paraburkholderia* species also have been implicated in plant health, and the *P. unamae* isolate which we found to confer resistance to phenazines in this study was originally isolated from the roots of corn ^39^. Understanding how often protective bacterial-fungal interactions occur in the rhizosphere may aid efforts to predict which microbial community compositions impact crop yield, differential stress tolerance of crops, and susceptibility to invasive pathogens. Additionally, though relatively little is known about phenazine interaction with the more deeply penetrating arbuscular mycorrhizal fungi that help support tree survival, a recent study suggests that phenazines could become toxic to this group of fungi if phenazine assault co-occurs with other environmental stresses ^48^. Finally, soil and plant-associated fungi are known to play a key role in carbon sequestration, indicating that understanding how fungi can be included into phenazine replete environments matters not just for plant growth on a warmer Earth, but also for maximizing our natural reservoirs of sequestered carbon even as soil becomes less able to sequester carbon as global temperatures rise ^3,49^.

Importantly, it is also possible that fungal-bacterial partnerships are exploited by fungal pathogens as well, as demonstrated by *P. edwinii* being capable of protecting a pathogenic *Fusarium* isolate in this study. One study showed that when *Paraburkholderia glathei* was paired with the fungal plant pathogens *Alternaria alternata* and *Fusarium solani*, *P. glathei* upregulated protein expression associated with antibiotic tolerance and oxidative stress response, while downregulating its starvation response ^50^. These results raise the tantalizing possibility that this may be another example of a mutually beneficial interspecies interaction competent to resist phenazine assault, where the bacterial stress responses may serve to protect its partner fungus from other bacteria rather than defending itself from its fungal host. Whether similar dynamics can play out in the context of the human host is also worth exploring, given that pathogenic fungi such as *Aspergillus fumigatus* and *Candida albicans* can be co-isolated from the lungs of cystic fibrosis patients with *Pseudomonas aeruginosa* ^51,52^. Lung function is negatively correlated with such coinfections, yet these fungi are largely inhibited by the phenazines produced by *P. aeruginosa* ^32,53^. Whether protective bacterial partners might help resolve this paradox remains to be seen.

Given the ease with which we isolated our model *P. edwinii-Aspergillus* pairing and the relative promiscuity of the protective bacterial partner and/or fungus being protected, we posit that interspecies cooperation may be an important method by which fungal membership in microbial communities is determined. This insight has important implications for diverse problems concerning environmental and human health. We hope that the model system established in this study will enable basic biological insights to be gained that will facilitate such partnerships to be exploited for human benefit in the future.

## Materials and Methods

### Strains and Media

Strains used in this study are listed in Table S2. *E coli* S17 was used for cloning and conjugation of the pMQ30 suicide vector during construction of deletion mutants. E. coli strain B2155 was used for mating of the mini-mariner containing pSC189. All *E. coli* were grown overnight in 5 mL lysogeny broth (LB), supplemented with 300 μM diaminopimelic acid (DAP) for B2155. Strains were grown shaking at 250 RPM at 37 °C for cloning constructs and 30 °C when used for conjugations. *Pseudomonas fluorescens* 2-79, *Pseudomonas chlororaphis*, *Pseudomonas chlororaphis* subsp. *aureofaciens* were grown overnight in 5 ml LB at 30 °C shaking at 250 rpm. *Paraburkholderia phenazinium*, *Paraburkholderia unamae*, *Paraburkholderia SOS3*, and *Paraburkholderia edwinii* were grown 24 hours under the same conditions in 5 mL potato dextrose broth (PDB) unless otherwise indicated.

The environmental *Aspergillus* isolate was grown on potato dextrose agar (PDA) at 30 °C, and all experiments which paired *P. edwinii* and the *Aspergillus* isolate were grown for 48 hours unless otherwise indicated. Clinical *Aspergillus fumigatus* as well as *Penicillium* species isolates were obtained from Children’s’ Hospital Los Angeles, and were grown under the same conditions. Growth on PDA for 1 week at 30 °C to allow conidiation was performed on all fungi to collect spores for storage and experimentation. Fungal conidiospores were collected by scraping mature colonies with a pipette tip, filtering the spores through cheesecloth, and freezing in 15% glycerol. All PDA plates contained 1.75% agar, while all other plates contained 1.5% agar.

### Co-isolation of phenazine tolerant fungal-bacterial pairings

100 mg of material was collected within the top 3 centimeters of top soil from outside of the Beckman Institute on the campus of the California Institute of Technology (34°8’21.15’’N 118°7’36.05’’W). The collected soil was washed in 0.1% TWEEN® 20 and pulsed in a sonicator bath for one minute to break up larger soil components. A serial dilution of the suspension was plated to extinction on potato dextrose medium. Fungal colonies were screened for adherent bacterial associates via amplification of the 16S rRNA encoding region of the bacterial genomes, and discovered pairings were subjected to challenge on potato dextrose agar supplemented with 300 μM phenazine-1-carboxylic acid. Surviving co-colonies were serially passaged in yeast-peptone (YP) medium to isolate the bacterium. The fungal partners were cured by growth on YP medium supplemented with 50 μg ml^−1^ gentamicin, repatched onto PDA and grown for one week at 30 °C until conidiation. Spores were collected and frozen in 12.5% glycerol.

### Phenazine Protection Assay

*P. edwinii* was grown shaking overnight in potato dextrose broth at 30 °C. Cultures were normalized to OD600 of 2.5, diluted 1:5, and mixed 1:1 with a suspension of ~ 4 x 10^7^ spores collected from the *Aspergillus* species. 6 ul of this mixture was spotted onto potato dextrose plates supplemented with 300 μM PCA and allowed to grow for 48 hours before measuring the co-colony diameter with the aid of Keyence digital microscope (VHX-600).

### Mutant transposon screen of *P. edwinii*

Transposon mutagenesis of *P. edwinii* was achieved using a mini-mariner transposon housed in the pSC189 vector and mutants were generated as follows. *P. edwinii* was grown overnight in potato dextrose broth, and the pSC189 vector-containing B2155 strain of *E. coli* was grown in LB with 50 μg ml^−1^ kanamycin and supplemented with 300 μM diaminopimelic acid (DAP). The *E. coli* was sub-cultured for 3-4 h until early log phase was achieved. 1 mL of each culture was pelleted and washed in their respective media before being resuspended in 100 μl of YP. The strains were mixed together and several 5 μl replicates were plated on YP plates overnight. Colonies were then scraped up and grown on YP plates lacking DAP and containing 30 μg ml^−1^ kanamycin to select for transposon insertions. Colonies were picked after two days and grown overnight in a 96 well plate in YP containing 60 μg ml^−1^ kanamycin. Mutants were mixed with spores of the Aspergillus species and grown on potato dextrose supplemented with PCA as above to screen for a dysregulated protection response. Mutants of interest had their transposition insertion mapped using arbitrary PCR.

### Construction of in-frame deletion and complementation strains in *P. edwinii*

In-frame deletions were constructed in *P. edwinii* using homologous recombination as previously described in *Pseudomonas* species with modification ^54^. ~1 KB regions upstream and downstream of the genomic region to be deleted were cloned into the pMQ30 suicide vector at the SmaI site, and the resulting constructs were electroporated into the S17 *E. coli* strain. Matings were conducted as described above, and the resulting mated colonies were scraped up and plated on potato dextrose containing 50 μg ml^−1^ gentamycin and 15 μg ml^−1^ chloramphenicol. Colonies were restreaked on selective plates, and finally patched onto YP plates amended with 7.5% (w/v) sucrose. Candidate colonies grown after 48 h were screened by polymerase chain reaction to identify those containing the desired in-frame deletions.

The pBBR1MCS-2 expression plasmid was used for complementation experiments. The gene of interest plus a 24 bp region upstream of the start codon were cloned into the plasmid and electroporated into *P. edwinii* using the following protocol. *P. edwinii* was grown overnight in potato dextrose broth. 4-5 mL of the culture was spun down and washed twice with 20% (w/v) sucrose at room temperature, before resuspending it in 100 μl of 20% (w/v) sucrose. 100 ng of the plasmid was added to 50 μL of the resuspended culture and electroporated using standard *E. coli* settings. Cells were allowed to recover for 2 h in YP at 30 °C before plating to YP plates containing 30 μg ml^−1^ kanamycin. Colonies were grown shaking overnight in potato dextrose broth with 60 μg ml^−1^ kanamycin before being used in growth assays without antibiotic selection as described above.

### Biofilm assay

V8 medium was produced by diluting V8 tomato juice 1:5 with ddH_2_O the same day of the experiment. Overnight *P. edwinii* cultures grown in potato dextrose broth were normalized to an OD600 of 2.5 and diluted 1:67 in the V8 medium and vortexed. 100 uL was pipetted per well into 96 well microtiter dishes as described for other systems ^31^. Biofilms were stored in a humidified microchamber and allowed to grow for 24 hours at 30 °C before being stained with 0.1% crystal violet for 20 minutes, and rinsed twice with ddH_2_0.

### Motility assay

*P. edwinii* strains were tested for motility using a swim assay. A modified M9 medium lacking added NaCl was made using 0.35% agar. Strains of *P. edwinii* were normalized to an OD600 of 2.5 and 100 μL was pipetted into a 96 well microtiter dish. A p10 pipette tip was dipped into one of these wells and subsequently plunged into the swim agar. Plates were incubated at 30 °C for 72 h.

### Phenazine sequestration assay

5 μl each of a spore suspension from the Aspergillus species and an overnight culture of *P. edwinii* normalized to OD600 2.5 were spotted 5 ul apart from each other on potato dextrose agar supplemented with 300 uM PCA and grown until the P. edwinii colonies developed a distinct yellow hue, approximately 48 h. Material from the P. edwinii colonies was then collected and resuspended in 250 uL Phosphate buffer saline solution (PBS). The samples were centrifuged and the supernatant was read for absorbance at 365 nm in a spectrophotometer and compared to a standard curve to derive PCA concentration. The pelleted fraction of the sample was dried and weighed to normalize sequestered PCA to bacterial dry weight mass.

This procedure was repeated to determine the fraction of reduced PCA present, but samples were collected and analyzed in an anerobic chamber, and the reduced PCA was identified using an excitation of 365 nm and read at an emission of 528 nm on a BioTek plate reader. Concentration was calculated using a standard curve of reduced PCA and compared to the total PCA concentration to determine the reduced fraction.

### PCA Degradation Assay

*P. edwinii* was added to 5 ml of PDB that was spiked with 300 μM PCA. For combined bacterial-fungal samples, the *Aspergillus* isolate was pre-grown in the medium for 3 days to allow an appreciable amount of slower growing fungal biomass to coexist with the subsequently added *P. edwinii* and PCA. 250 μl was sampled every 24 hours for 3 days, and all cultures had reached stationary phase by the first sampling. Cells were pelleted, and the supernatant was used to quantify PCA by absorbance at 365 nm compared to a standard curve and PCA negative control.

### Microelectrode Profiling

*Aspergillus* and *P. edwinii* colonies were grown side by side or alone on PDA as above for 48 h at 30 °C. Unisense pH and redox 25 μm tip microelectrodes were paired with a steel reference probe (Unisense) in accordance with the manufacturer’s instructions for use and calibration, and were readthrough a high-impedance millivoltmeter-equipped multimeter, while a 10 μm tip O_2_ probe was read through the multimeter’s picoampere amplifier. Oxygen concentration was read at 10 μm interval depths within the colonies, while pH and redox values were read at 25 μm intervals through the colonies. Redox values are given relative to a standard hydrogen electrode. Initial calibration and recording of data were performed using the Unisense SensorTrace Suite software. All calibrations and measurements were conducted at 23 °C.

### MiPACT clearing and HCR

Fungal tissue was cleared using the MiPACT procedure as described previously with minor modification ^28^. Briefly, samples were grown for 48 h, and whole co-colonies cut from their growth medium, removing as much agar as possible. Samples were incubated at 4 °C overnight in 3% (v/v) paraformaldehyde. Samples were cleared for 2 weeks in a spinning 8% (w/v) SDS solution at 37 °C. Samples were washed in PBS, treated with 1 mg ml^−1^ lysozyme for 30 minutes at 30 °C before being hybridized with HCR 2.0 eubacterial probes and incubated overnight at 46 °C while gently shaking. Samples were washed and the amplification step was performed using hairpins tagged with an alexa 647 fluorophore for visualization. Samples were stained with 1 μg ml^−1^ DAPI, and microscopy was performed on an inverted confocal Leica model TCS SPE confocal microscope with a 10x objective for the colony center and edge, and a Nikon Ti2 Eclipse widefield microscope with a 4x objective for images of the colony ridge. Contrast of the HCR generated images were normalized to the brightest signal in like samples (i.e. colony edge vs edge, center vs center). Contrast was adjusted independently for each DAPI stained images for clarity of fungal morphology present near the bacteria.

### CFU Quantification

Fungal-bacterial co-colonies were grown as above, and the colonies were homogenized using a tissue homogenizer (Bio-Gen Pro200) at 60% power for one minute. The resulting slurry was serially diluted and plated to YP medium supplemented with 50 μg ml^−1^ nystatin to prevent fungal growth. Bacterial colonies were grown for 48 h and counted.

### HPLC-MS

*P. edwinii* colonies were scraped from their growth plates and resuspended in phosphate buffer saline before being pelleted. Supernatants were frozen and thawed to encourage precipitation of large particulate condiments, and centrifuged using a cellulose acetate Spin-X column (VWR). 20 ul of the supernatant was injected onto a Waters e2695 Separations Module equipped with a 2998 PDA Detector and run through a C18 column (XBridge, 3.5 um, 2.1 x 50 mm) housed at 40 °C at a flow rate of 0.5 ml min^−1^ for 20 mins. The mobile phase consisted of ddH20 + 0.04% (v/v) NH_4_OH with a gradient to 70% (v/v) acetonitrile + 0.04% (v/v) NH_4_OH with a constant background of 2% (v/v) methanol and compared against a prepared standard. The identify of PCA was confirmed using a quadrupole Time of Flight MS (Q-TOF, Xevo G2-XS, Waters) targeting a mass of 224.2 m/z.

### Genomic Sequencing and Annotation

*P. edwinii* genomic DNA was recovered using the Qiagen plant and tissue kit on 1 mL of culture grown overnight in PDB. Genomic DNA from the *Aspergillus* species was recovered by growing it for 72 h in PDB until large fungal aggregates had formed. These aggregates were frozen in liquid nitrogen, and crushed with a mortar and pestle followed by a chloroform extraction.

Illumina sequencing of both genomes was conducted at the Caltech genomic core facility, and subsequent PACBio sequencing was carried out at the UC Irvine Genomics High Throughput Facility. Genome assembly was performed using SPAdes version 3.12.0, and BASys was used for genome annotation of *P. edwinii* as provided by the Health Sciences Library System at the University of Pittsburg.

## Supporting information

SI Legends, Ref and Figs. S1-S4

## Acknowledgements

We thank members of the Newman lab for constructive feedback on the project and the manuscript, and The Millard and Muriel Jacobs Genetics and Genomics Laboratory at Caltech and Igor Antoshechkin for support during library preparation and sequencing. We thank Marko Kojic for help screening transposon mutants, as well as Robert Cramer, Deborah Hogan, and Jeff Holloman for sharing their expertise in mycology. This work was supported by the Life Sciences Research Foundation (postdoctoral fellowship to K.M.D.), the Resnick Institute (K.M.D. and D.K.N.) and the NIH (1R01AI127850-01A1 to D.K.N.).

