## Supplementary material for "*Paraburkholderia edwinii* protects *Aspergillus* sp. from phenazines by acting as a toxin sponge": SI Legends, Ref and Figs. S1-S4

### Supplemental Figures

**Fig. S1**. **Fungi from identified bacterial-fungal pairings are sensitive to PCA.**. **A)** *Lecythophora* and *Aspergillus* species grown on PDA, PDA supplemented with 300 µM PCA, and finally in the presence of PCA along with co-isolated bacterial partners, a *Paraburkholderia* species and a *Luteibacter* species, respectively. Plates were grown for 48 hrs at 30 ºC. **B)** Phenazine production is required to inhibit growth of the *Aspergillus* species. *Pseudomonas chlororaphis* and *Pseudomonas fluorescens* inhibit fungal growth, but the *::phzD* and *::phzB* strains, respectively, fail to inhibit fungal growth after growing on PDA for 48 hrs at 30 ºC.

**Fig. S2**. **Bacterial aggregates accumulate along the ridge of co-colonies treated with PCA.** A band of bacteria can be seen in both treated and control colonies between the inner and outer zones. In the PCA treated colony, this band is thicker with more well formed aggregates, and is representative of the region larger aggregates are visible to the eye during co-colony growth on PCA. Samples were cleared using the MiPACT protocol, and images were captured on a Nikon Ti2 Eclipse. Red represents the signal produced by eubacterial HCR probes coupled to Alexa fluor 647 nm..

**Fig. S3**. ***P. edwinii*** **does not degrade PCA in liquid culture**. *P. edwinii* was grown in potato dextrose broth shaking at 30 ºC spiked with and without 300 µM PCA, and with PCA and its fungal partner. PCA concentration was measured every 24 hours for three days. A control lacking *P. edwinii* was included to confirm PCA did not degrade over this time period. Quantification occurred by measuring absorbance at 365 nm. Error bars represent standard deviation of four biological replicates.

**Fig. S4**. **Morphology of WT and Δ*hrcA* P. edwinii and properties of the *Aspergillu*s isolate with and without PCA challenge**. **A)** Comparison of the *Aspergillus* isolate grown alone (left, top), with WT *P. edwinii* (center), and with the Δ*hrcA* mutant (bottom). Note large aggregates in the Δ*hrcA* mutant despite the absence of PCA **B)** WT (left) the the Δ*hrcA* mutant (right) grown in the absence (top) and presence (center) of PCA, and next to its partner fungus when challenged with PCA (bottom). The WT strain shows a thickening of the colony in response to PCA, as well as an increase sheen. A similar thickening and sheen is visible for the Δ*hrcA* mutant in all conditions. **C)** The *Aspergillus* isolate produces an oxidizing environment when growing on potato dextrose agar in the absence of PCA, but generates a more reducing environment during challenge with 300 µM PCA. Error bars represent the standard deviation of 3 measurements at each depth. All colonies were grown for 48 hours at 30 ºC.

### Supplemental Tables

**Supplementary Table 1.** List of transposon mutants found in P. edwinii that alter its ability to protect its partner fungus, including phenotype (left), closest homologue of P. edwinii gene each transposon hit was found in based on NCBI BLAST using arbitrary PCR on the mutant strains, and any apparent homologue the gene encodes.

**Supplementary Table 2.** List of strains and plasmids used in this study.

Figure S1

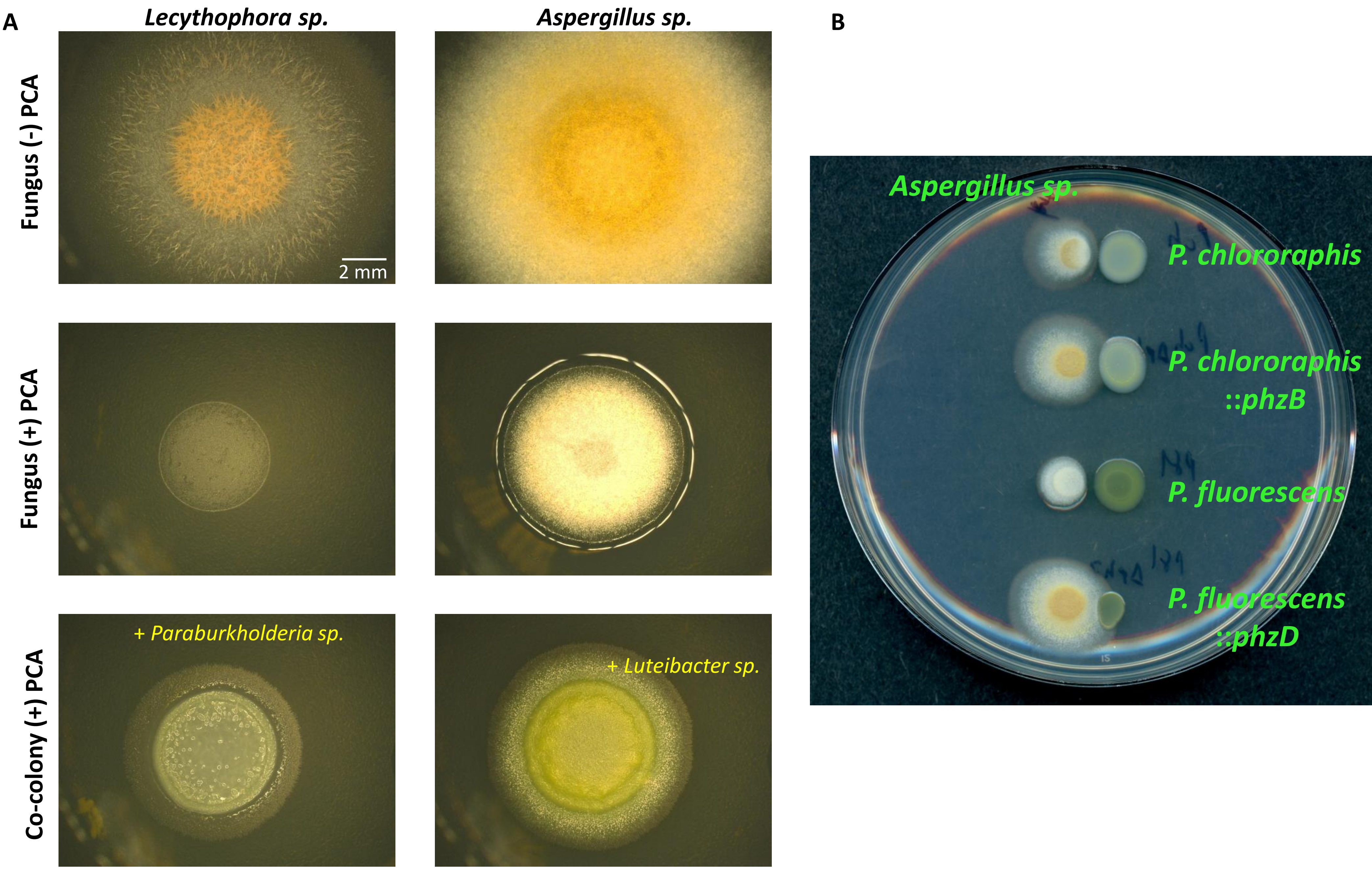
A
Lecythophora sp.
Aspergillus sp.
Fungus (-) PCA
Fungus (+) PCA
Co-colony (+) PCA
2 mm
+ Paraburkholderia sp.
+ Luteibacter sp.
B
Aspergillus sp.
P. chlororaphis
P. chlororaphis
::phzB
P. fluorescens
P. fluorescens
::phzD

Figure S2

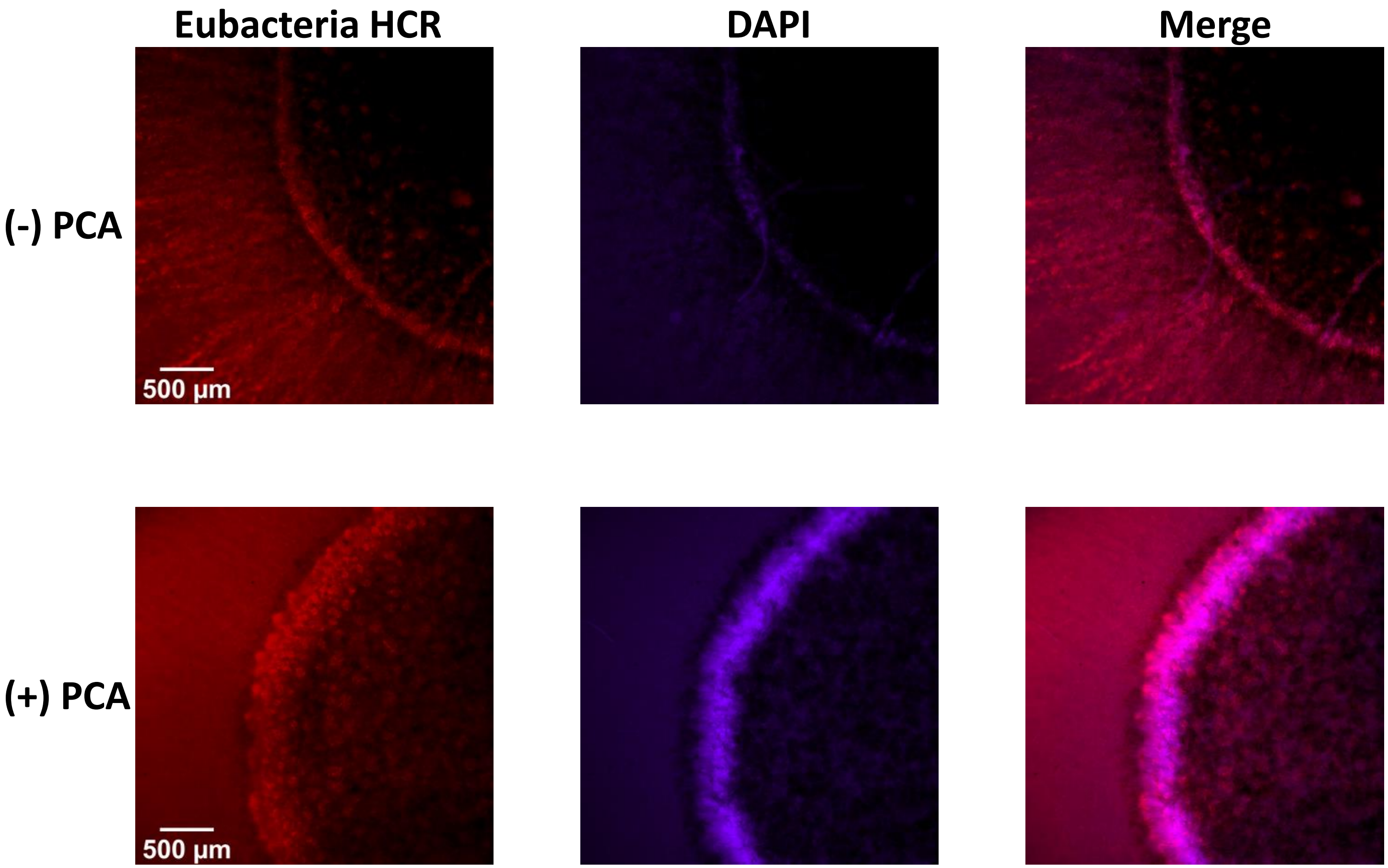

**Figure S3**

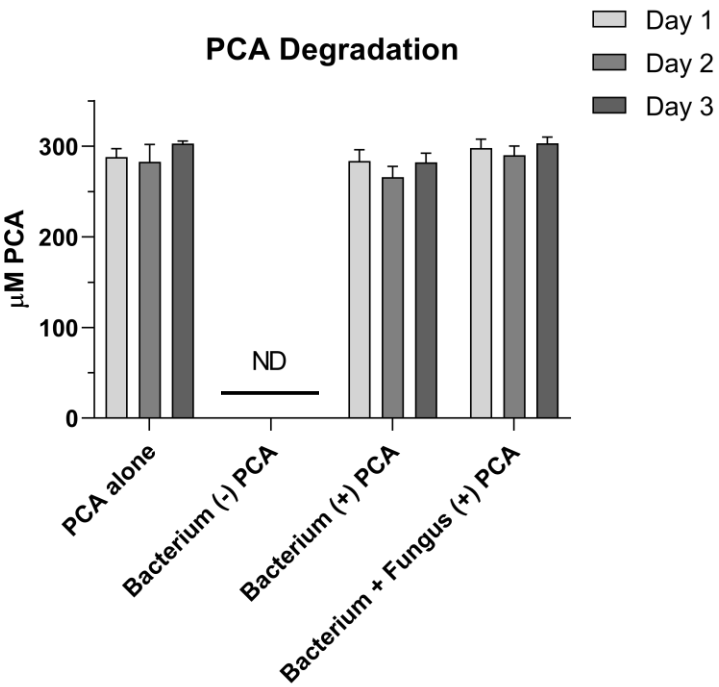
PCA Degradation
Day 1
Day 2
Day 3
µM PCA
300
200
100
0
ND
PCA alone
Bacterium (-) PCA
Bacterium (+) PCA
Bacterium + Fungus (+) PCA

Figure S4

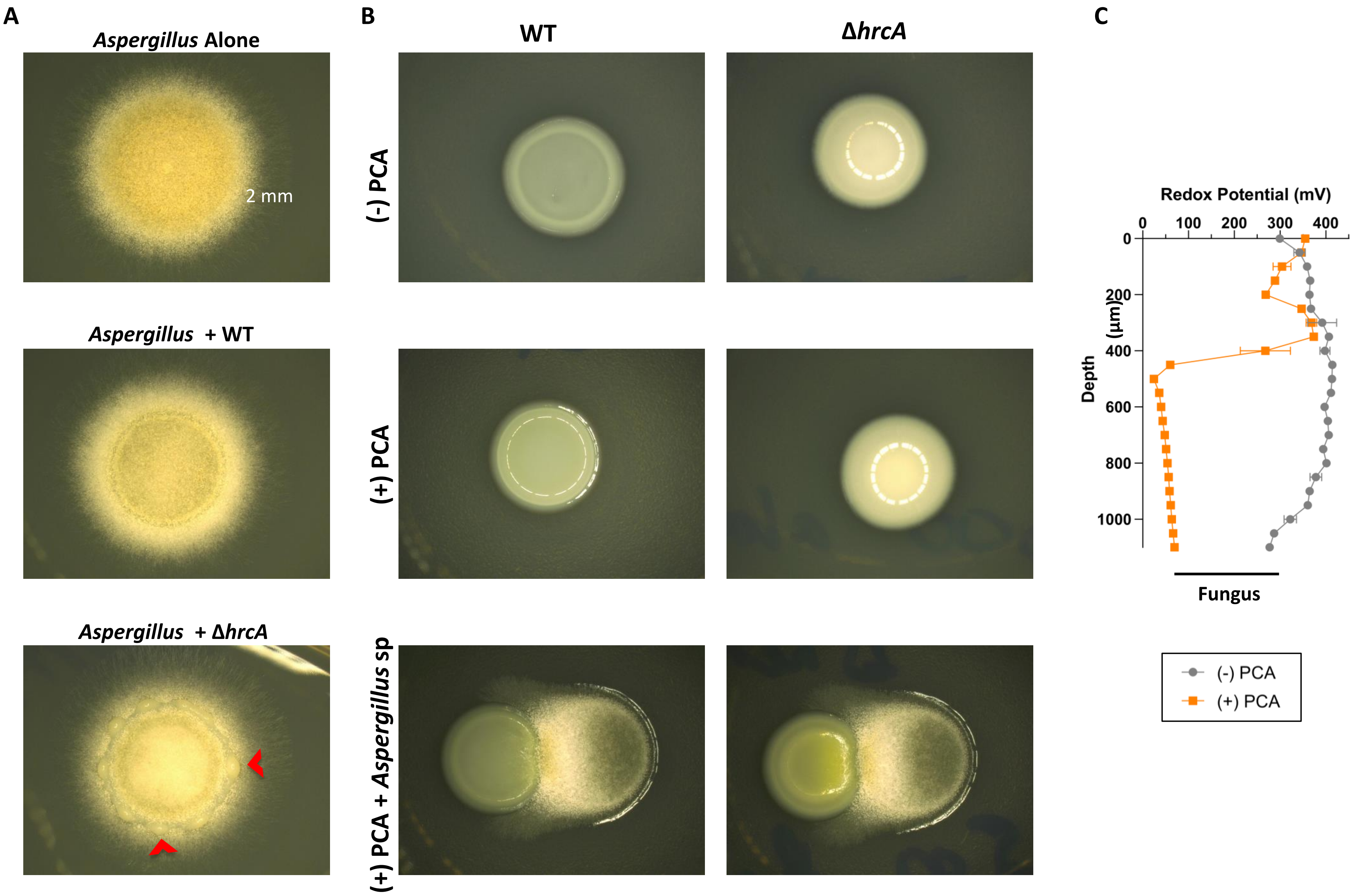

A
Aspergillus Alone
2 mm
Aspergillus + WT
Aspergillus + ΔhrcA
B
WT
ΔhrcA
(-) PCA
(+) PCA
(+) PCA + Aspergillus sp
C
Redox Potential (mV)
0
100
200
300
400
Depth (µm)
0
200
400
600
800
1000
Fungus
(-) PCA
(+) PCA
